# Measuring the average power of neural oscillations

**DOI:** 10.1101/441626

**Authors:** Liz Izhikevich, Richard Gao, Erik Peterson, Bradley Voytek

**Author notes:** Correspondence should be addressed to Bradley Voytek.

## Abstract

**Background:** Neural oscillations are often quantified as average power relative to a cognitive, perceptual, and/or behavioral task. This is commonly done using Fourier-based techniques, such as Welch’s method for estimating the power spectral density, and/or by estimating narrowband oscillatory power across trials, conditions, and/or groups. The core assumption underlying these approaches is that the mean is an appropriate measure of central tendency. Despite the importance of this assumption, it has not been rigorously tested.

**New method:** We introduce extensions of common approaches that are better suited for the physiological reality of how neural oscillations often manifest: as nonstationary, high-power bursts, rather than sustained rhythms. Log-transforming, or taking the median power, significantly reduces erroneously inflated power estimates.

**Results:** Analyzing 101 participants’ worth of human electrophysiology, totaling 3,560 channels and over 40 hours data, we show that, in all cases examined, spectral power is not Gaussian distributed. This is true even when oscillations are prominent and sustained, such as visual cortical alpha. Power across time, at every frequency, is characterized by a substantial long tail, which implies that estimates of average power are skewed toward large, infrequent high-power oscillatory bursts.

**Comparison with existing methods:** In a simulated event-related experiment we show how introducing just a few high-power oscillatory bursts, as seen in real data, can, perhaps erroneously, cause significant differences between conditions using traditional methods. These erroneous effects are substantially reduced with our new methods.

**Conclusions:** These results call into question the validity of common statistical practices in neural oscillation research.

**Highlights:**

- Analyses of oscillatory power often assume power is normally distributed.
- Analyzing >40 hours of human M/EEG and ECoG, we show that in all cases it is not.
- This effect is demonstrated in simple simulation of an event-related task.
- Overinflated power estimates are reduced via log-transformation or median power.

## Intro

Spectral analysis is an indispensable tool for many cognitive, systems, and clinical neuroscientists. It is used to characterize oscillations in meso- and macro-scale field potential data ranging from invasive local field potential (LFP) and electrocorticography (ECoG) to non-invasive electro- and magnetoencephalography (EEG and MEG). In particular, fixed bands of oscillatory power are often used to predict behavioral or disease conditions. For example, delta (1-4 Hz) power correlates with different stages of sleep (Amzica and Steriade, 1998); alpha (8-12 Hz) power decreases topographically during visual attention (Palva and Palva, 2007; Voytek et al., 2017); beta (15-30 Hz) power is pathologically strong during the progression of Parkinson’s disease (Alonso-Frech, 2006; McCarthy et al., 2011). Power in these bands can be assessed via the power spectral density (PSD), which is typically calculated using Welch’s method (Voytek et al., 2010), which estimates the power at any frequency as the arithmetic mean of many windowed Fourier decompositions across time, or across experimental trials of the same condition (Welch, 1967). A related approach is the event-related spectral response, where spectral power at each frequency is averaged across trials for each time point, which contains not one, but two spectral power estimates based on the arithmetic mean: 1) across trials, and, 2) across time (Makeig, 1993). More sophisticated methods such as wavelet filtering or multitaper analysis use different windowing methods and convolution kernels (Bruns, 2004), but the final step frequently involves averaging over time. These averages are then statistically compared across different experimental conditions to establish whether there is a significant power difference between them.

The methods described above are used in nearly every study looking at oscillatory power changes. Welch’s method has a long history in science and engineering applications, with nearly 7000 citations, according to Google Scholar, at the time of this writing. An important assumption of Welch’s method, and averaging in the frequency domain in general, is that the noise at any single frequency is symmetrically or Gaussian distributed and stationary. This assumption must be held for the arithmetic mean to be a reliable measure of central tendency. If this assumption is violated, the estimate is severely skewed by relatively few outliers.

There are strong hints in the existing literature that power is not Gaussian distributed. First, spectral power follows a 1/f (power law-like) distribution across frequencies (Freeman and Zhai, 2009; Miller et al., 2009; Baranauskas et al., 2012; He, 2014; Podvalny et al., 2015; Gao, 2016; Gao et al., 2017; Haller et al., 2018) despite recording modality (Voytek et al., 2010, 2015). Second, from a physiological perspective, it is clear that firing rate, dendritic connections, synaptic weights, and other parameters are distributed according to the log-normal, gamma, or other long-tailed distributions (Buzsáki and Mizuseki, 2014). Given that these are the parameters responsible for generating LFPs, and that the power spectrum is 1/f, we hypothesize that oscillatory power is long-tailed, rather than symmetric or Gaussian. The assumption that spectral power of neurophysiological signals is Gaussian distributed across time has never been explicitly examined. Testing this is critical, as any study in which the arithmetic mean is used will have a poor estimate of the true central tendency. To demonstrate by way of a simple analogy: if there are 9 people in a room with an average net worth of $50,000, and a billionaire walks into the room, it does not seem terribly “average” to say that the average net worth of the 10 people in the room is now $100,045,000.

In this study, we examined the distribution of spectral power in a large range of human electrophysiology recordings (EEG/MEG/ECoG). To be as comprehensive as possible, the datasets we analyze are heterogeneous in terms of sampling rate, experimental condition, participant age, and various other parameters (**Table 1**). We first show that the distribution of neural recording power *across time*, in every frequency bin, is significantly skewed, non-Gaussian, and better approximated by a gamma distribution across all participants and recording modalities (**Fig. 2**). We then show, within participants, that this skewness holds true for all modalities, and across all brain regions (**Fig. 3**). Finally, we show, in simulation, how averaging power across trials, when power in those trials are gamma-distributed, can lead to inflated significance (**Fig. 4**).

**Table 1.**
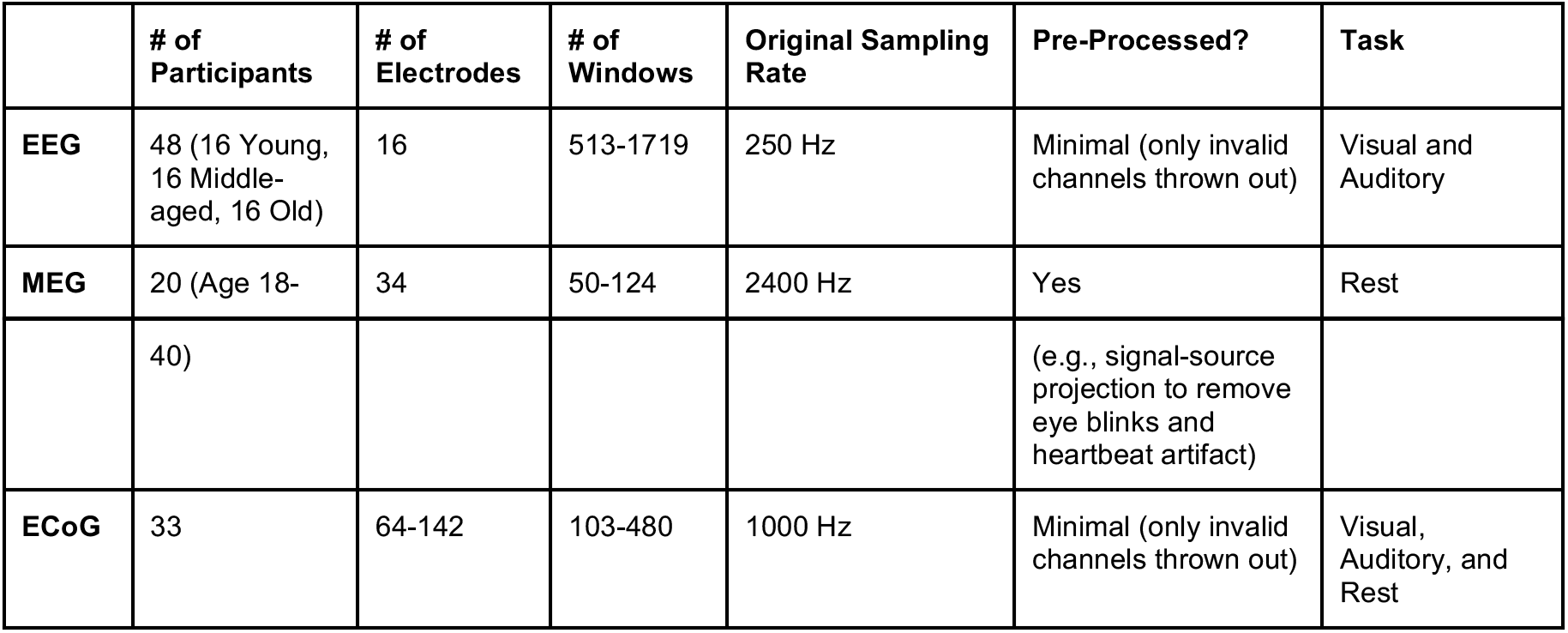
Description of analyzed datasets.

|  | # of Participants | # of Electrodes | # of Windows | Original Sampling Rate | Pre-Processed? | Task |
| --- | --- | --- | --- | --- | --- | --- |
| EEG | 48 (16 Young, 16 Middle-aged, 16 Old) | 16 | 513-1719 | 250 Hz | Minimal (only invalid channels thrown out) | Visual and Auditory |
| MEG | 20 (Age 18- | 34 | 50-124 | 2400 Hz | Yes | Rest |
|  | 40) |  |  |  | (e.g., signal-source projection to remove eye blinks and heartbeat artifact) |  |
| ECoG | 33 | 64-142 | 103-480 | 1000 Hz | Minimal (only invalid channels thrown out) | Visual, Auditory, and Rest |

**Figure 1.**
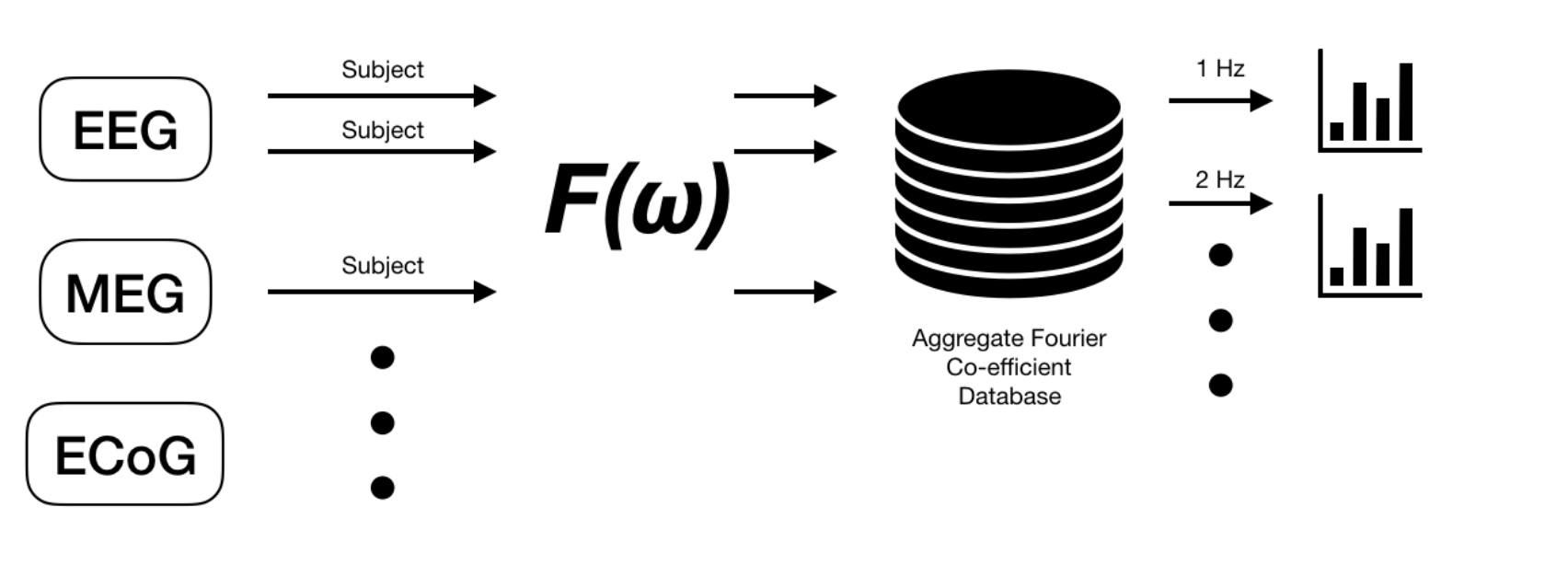
Analysis pipeline. EEG, MEG, and ECoG time series from multiple participants and electrodes are processed via short-time window Fourier transform (*F(ω)*). Squared magnitude of the Fourier coefficients are stored in a database of power values, from which we query by aggregating across different dimensions (e.g., participant or electrode). The resulting aggregated histograms represent distribution of power in neural recordings at frequencies up to 25Hz, exemplified in Figures 2 and 3, which are subsequently fit with gamma distributions to measure skewness.

**Figure 2.**
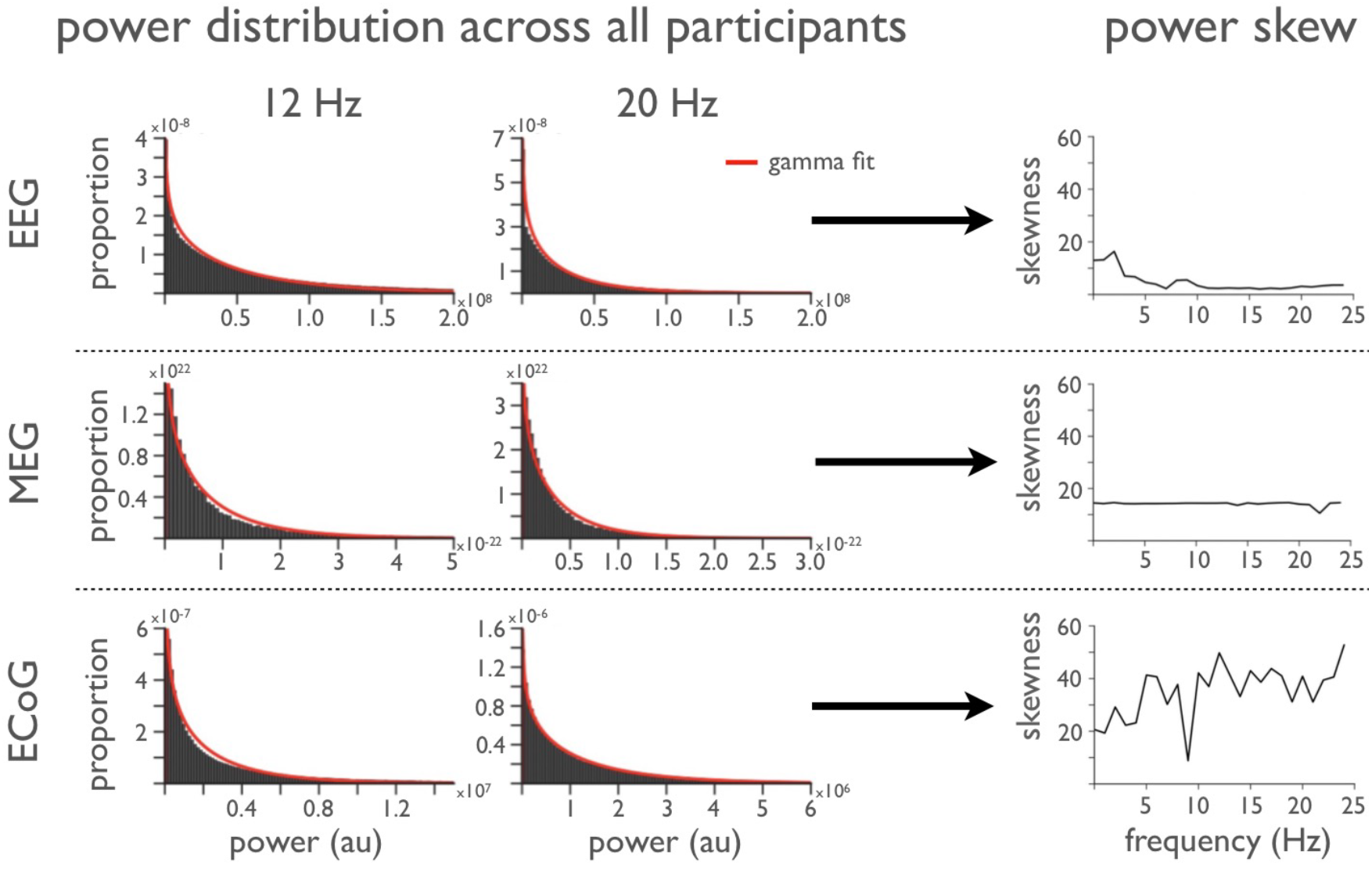
Narrowband power distributions (12 and 20 Hz) for all participants, electrodes, and time windows. Canonical alpha (12 Hz; first column) and beta (20 Hz; second column) band power in recordings of different modalities all show heavily skewed distributions when aggregated across electrodes and participants. Skewness, measured as the shape parameter from gamma distribution fits, is non-zero for all frequencies up to 25 Hz (third column).

**Figure 3.**
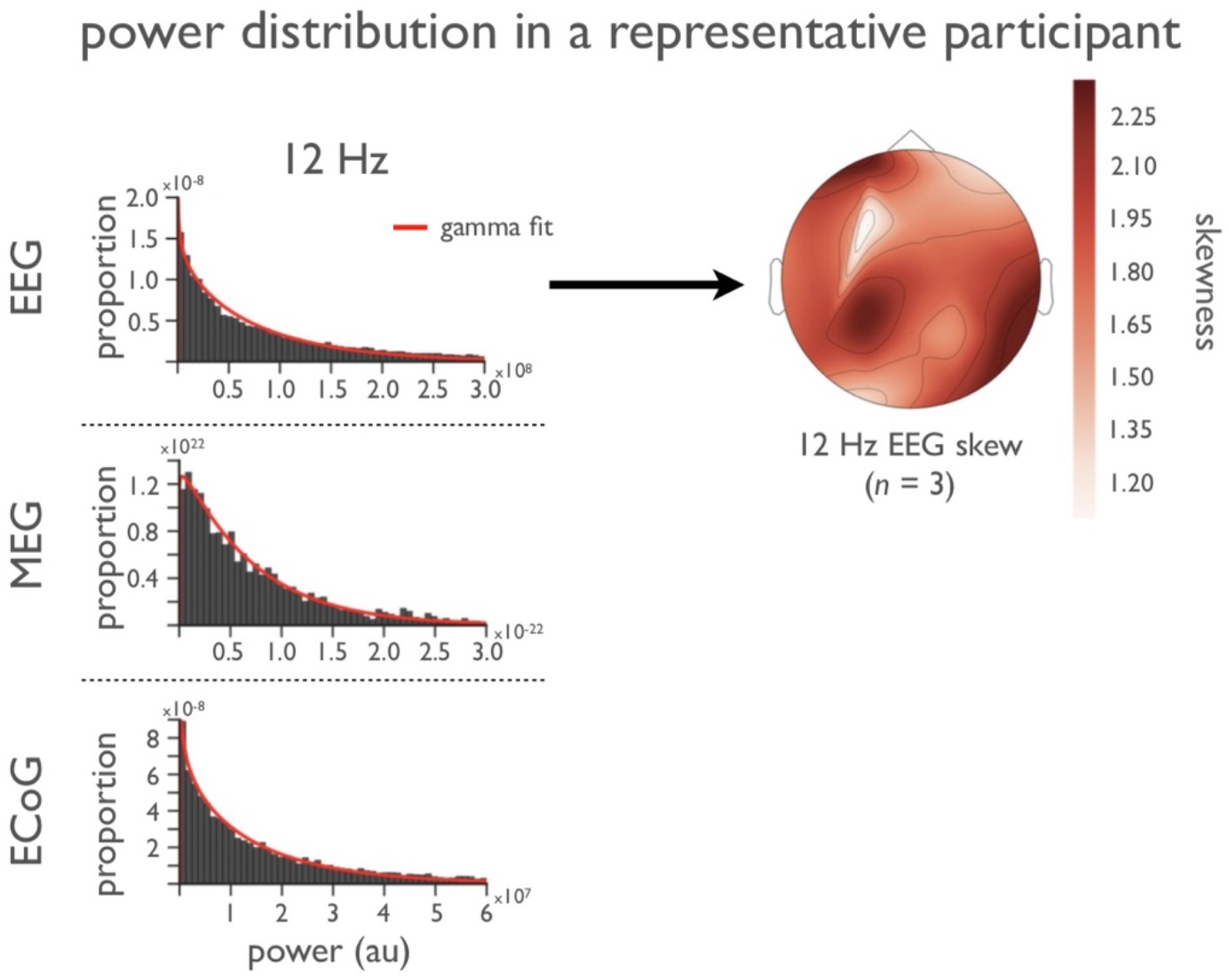
Example narrowband power distributions (12 Hz) across one individual participant. Power at 12 Hz in all 3 modalities follow skewed distributions, and skewness is non-zero in all EEG electrodes across the scalp for the plotted representative participant. Note the skewness is always above 1.0, far from the 0.0 of a Gaussian distribution.

**Figure 4.**
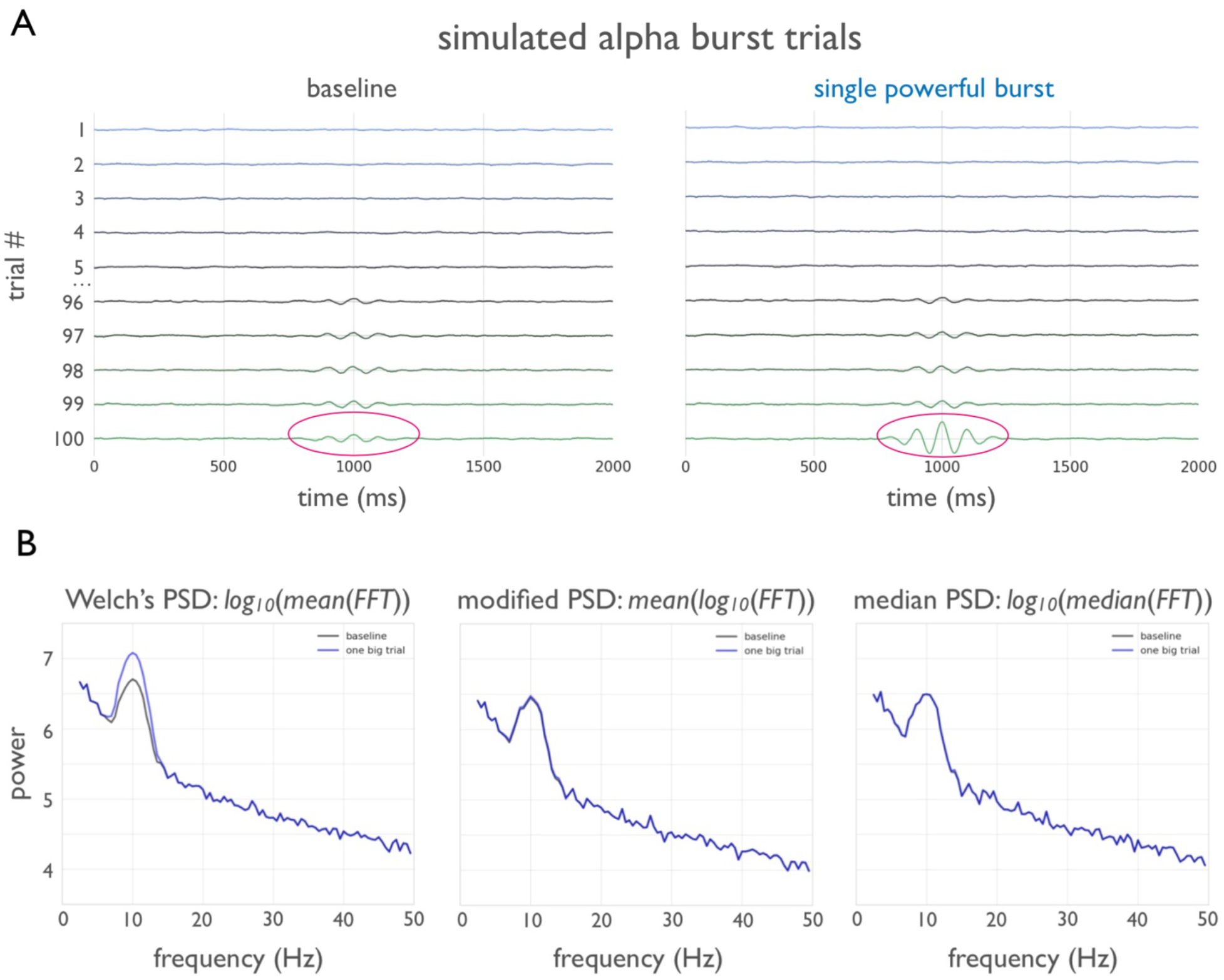
Effects of a single large-amplitude oscillatory event on Welch’s PSD. **A)** When the alpha oscillation amplitude is quadrupled in the single trial with the highest power, out of 100 simulated trials, Welch’s method (arithmetic mean, left) shows a drastic 10 Hz power difference between the two simulated sets of trials. **B)** In contrast, log-transforming first and then taking the mean (middle), or taking the median (right), reduces the undue influence of this rare, high-powered oscillation.

These results show that averaging oscillatory power estimates, using common methods, results in poor estimates of central tendency that are significantly inflated by the rare, high-power events that dominate electrophysiological recordings. That is, our results call into question the validity of any study which attribute their results to changes in the PSD as measured using Welch’s method and/or the arithmetic mean of trial or group-averaged spectral profiles. We conclude with several possible recommendations for future studies examining average spectral power.

## Methods and Data

### Data

To study the distribution of spectral power in multiple recording modalities, we first create a database of power spectra using the data specified in **Table 1**. EEG data is provided by the Kutas Lab at UC San Diego (unpublished); MEG data are CTF MEG from the OMEGA open database (Niso et al., 2016), and ECoG data are from Johns Hopkins and UC San Francisco. For the MEG data, every eighth MEG sensor was chosen to reduce total data size, giving a final set of 34 channels per subject. All participants gave informed consent approved by the Institutional Review Boards at the respective recording institutions.

We examine the distribution of spectral power at all frequencies 1-25 Hz and report on three specific frequencies—6, 12, 20 Hz—corresponding to three common, canonical oscillatory bands of theta, alpha, and beta respectively.

### Fourier Power Distributions

The database-creation workflow and analysis pipeline are outlined in **Fig. 1**. First, all data is down-sampled to 250 Hz. Next, each channel, for each participant, is broken up into non-overlapping 5-second windows and Fourier transformed. Then all Fourier coefficients are aggregated, for each channel and participant, resulting in a distribution of power values for each frequency. Distribution of power, at specific frequencies, is then analyzed across all participants, in aggregate (**Fig. 2**) and individually (**Fig. 3**).

### Simulation

We simulate 100 trials of data, where each trial has a brown noise background signal (*f^-2^*) of equal power. Brown noise is simulated by convolving a sequence of numbers drawn from a normal distribution with an exponentially decaying filter. The oscillatory component is simulated as a Gaussian-modulated sinusoid, with squared amplitudes drawn from an exponential distribution to match empirical data, which is added to the brown noise signal. Finally, the 100 trials are sorted by ascending oscillatory power, and the 5 lowest and 5 highest powered trials are plotted in **Fig. 4A** (left, baseline). In the ‘burst’ conditions (**Fig. 4A**, right), the oscillatory signal of the trial with the highest amplitude is multiplied by 4, simulating a rare and high-amplitude signal; the rest of the trials are otherwise kept identical. Subsequently, we perform the above replacement procedure for the *N* highest-powered trials to determine the minimum number of trials required to produce a statistically significant difference under Student’s *t*-test (p<0.05). Note that the assumptions of normality for the t-test are explicitly broken in these simulated data, emulating the violations we demonstrate in real data and emphasizing how these violations cause inflated significance.

## Results

The distribution of power values across time, across participants, and across electrodes, are never Gaussian distributed, regardless of recoding modality (EEG, MEG, or ECoG). While the detailed analyses below focus on the common EEG and MEG oscillatory range (1-25 Hz), the same non-Gaussian skew is observed in all frequencies up to our maximum analysis range of 100 Hz.

### Aggregate analysis of power distribution

We report that, when aggregating the spectral power of each frequency from all analysis windows, electrodes, and participants per data modality, spectral power does not follow a Gaussian distribution. Rather, the distribution of power for each frequency is well fit by a gamma distribution (**Fig. 2**). The gamma distribution is described by two parameters—shape and rate—where the shape parameter is proportional to the overall skewness of the distribution while the rate parameter describes the dispersion, or spread, of the data. For a Gaussian distribution, the shape parameter is 0.0.

We estimate the gamma parameters of the aggregated data at each frequency between 1-25 Hz. As seen in **Fig. 2**, none of the shape parameters are near zero (the skew of a Gaussian distribution). Rather, the skew is always positive, with a heavy tail caused by rare, large power events that pull the mean away from the median and mode.

### Analysis of power distribution across participants

To confirm that the skewness of the aggregate spectral power is not due to the skewness of a particular participant’s data or from aggregating across participants, we analyze the distribution of spectral power per individual participant, while still aggregating spectral power of all electrodes and time windows for that particular participant. The skewed distributions are observed across all participants. The distribution of spectral power is heavily skewed for each individual participant at all frequencies, and also at all frequencies at every electrode. **Fig. 3** illustrates a heavy skew in the 12 Hz frequency band.

### Analysis of power distribution across space

To confirm that the skewness of spectral power is non-zero across space, we also analyze the distribution of spectral power per electrode in EEG recordings, while still aggregating across all participants and time windows for the particular electrode. Across all electrodes, a constant positive skew is still observed (**Fig. 3**). The skewness of spectral power at 12 Hz never reaches 0, no matter where the electrode on the scalp is located. Furthermore, as seen in the left of **Fig. 3**, the mean skewness never reaches near zero at any frequency or electrode combination.

### Effect of rare, high power oscillatory events on power averages

To demonstrate the effect of rare, high power oscillatory events on significance tests between simulated conditions, we simulate brown noise (*f^-^*^2^)-distributed time series with 10 Hz oscillations of gamma-distributed power (see Methods). We simulate 100 trials using this procedure, and copy those 100 trails, producing two identical sets of simulated trials. Because of the gamma distribution of power, most oscillations are very weak, with a few that are quite powerful (**Fig. 4A**).

We then artificially increase the amplitude of a single oscillation. Specifically, the highest amplitude trial is multiplied by a factor of 4 to simulate a rare, but not implausible, high-amplitude event, in one of the two previously-identical sets of conditions. Note that the 99 other trials in the two sets remain identical. Doing so dramatically increases the average amplitude in the oscillatory band when the PSD is estimated as the *log_10_(mean(FFT))* of the time series, as per Welch’s method (**Fig. 4B**, left).

We then repeat this procedure, multiplying the next-highest amplitude trial by a factor of 4. Each time we increase the amplitude of a single trial, we compute a Students t-test of significance between the two simulated sets of 100 trials, comparing power at 10 Hz. We find that swapping just 5 of the 100 trials, using this procedure, results in a statistically significant difference under Student’s t-test (*p*<0.05).

Modifying this slightly to mean*(log_10_ (FFT))* or *log_10_(median(FFT))* largely mitigates this issue (**Fig. 4B**).

## Discussion

### Implications of non-Gaussian distribution of oscillations

Our results demonstrate that human EEG, MEG, and ECoG data manifest skewed power distributions, which result in biased central tendency estimates when the mean is used. Our results imply an inherent statistical flaw in studies where average power is computed across time, either from Fourier-based power estimates like above or from convolution-based estimates via filtering. Subsequent hypothesis-testing which compares average spectral power estimates using common procedures such as *t*-tests, ANOVA, and other similar null hypothesis statistical significance tests that assume Gaussianity, will therefore be flawed. As we demonstrated via simulation, increasing the power of just a few simulated trials produces a significant increase in estimated mean power. Whether such a rare event—just a few outlier power spectral events—are meaningful, from a physiological perspective, is difficult to determine.

### Addressing caveats

Throughout our analysis we see that electrophysiological activity is not Gaussian distributed. Nonetheless, we keep in mind the following caveats. First, it is undeniable that, though we are working with over 100 participants worth of data across three different electrophysiological recording modalities, including both task and non-task recordings, there could exist a particular recording modality or a particular group of participants that exhibit more Gaussian-distributed spectral power. However, given the heterogeneity of our data and the consistency across data types, it is likely that oscillatory power is not Gaussian distributed. We report that analyses of non-human primate LFP data showed similar non-Gaussian distributions of electrophysiological power.

Second, as seen in **Table 1,** the data we analyzed were not all consistently pre-processed. The EEG and ECoG data only had minimal pre-processing where only invalid channels of recording were thrown out. The MEG were also pre-processed, yet we cannot guarantee that *all* non-neural artifacts were removed. It is possible that the pre-processing reduces the variance of skew, as seen in **Fig. 2,** when comparing the distributions of skews of MEG to EEG and ECoG. Regardless, in all cases that we have seen so far, we find that an extremely heavy skew is present. In no cases is skew near zero. Therefore, we argue that our results are robust: common variants of electrophysiological data do not have a Gaussian distribution.

### Alternatives to Welch’s Method

Methods that assume the arithmetic mean to be a valid measure of central tendency should be used only after an appropriate data transform is performed prior to averaging, *e.g.*, computing the mean on log-transformed power (Smulders et al., 2018), as illustrated in **Fig. 4B**. Alternatively, one can use a “Median Welch” method, where the median power at each frequency is taken, rather than the mean. In comparing traditional Welch’s method to both log-transformed and median on simulated data (**Fig. 4**), we show how Welch’s method is biased by the long-tail power values present in the data.

Concretely, the spectral power using Welch’s method increases significantly, even when just a few trials of data are amplified. This is because Welch’s method inherently tries to average a squared value (power is proportional to amplitude squared). Therefore, unless the intended effect is to intentionally amplify the effect of rare oscillatory bursts—which may be a desirable trait—we recommend that arithmetic mean-based averaging of spectral power is avoided. We recommend alternatives, such a log-transformed or median approaches, that are more resilient against the effects of rare events.

A strong implication of our observation, wherein power at all frequencies in electrophysiological data is long-tail distributed, is that data are at most times very low power and only in very rare instances do high power bursts occur. This is important because many oscillation analysis methods also assume stationarity. However, from a cognitive and behavioral standpoint, this is particularly interesting given that emerging work is highlighting the potential functional role of oscillatory bursts (Feingold et al., 2015; Lundqvist et al., 2016), as opposed to sustained oscillations (Peterson and Voytek, 2017). Additionally, because common frequency-domain methods conflate higher power oscillations with longer-lasting oscillations (Jones, 2016), it may be more appropriate to complement frequency-domain analyses with time-domain approaches that can discriminate between those differences (Cole and Voytek, 2018).

## Acknowledgements

We thank Thomas Donoghue and Priyadarshini Sebastian for their assistance with data preparation and comments on a draft of this manuscript. Izhikevich is supported by the National Science Foundation Graduate Research Fellowship Program (DGE-1656518) and a Stanford Graduate Fellowship. Gao is supported by the Natural Sciences and Engineering Research Council of Canada (NSERC PGS-D), UCSD Kavli Innovative Research Grant (IRG), and the Katzin Prize. Voytek is supported by a Sloan Research Fellowship (FG-2015-66057), the Whitehall Foundation (2017-12-73), and the National Science Foundation under grant BCS-1736028. The authors declare no competing financial interests.

**Author contributions**
All authors initiated and designed the study. Izhikevich built the database and analysis pipeline, and all authors analyzed the data. All authors contributed to the manuscript.

